# IntraTalker - Modelling Intracellular Signalling in Cellular Crosstalk

**DOI:** 10.64898/2026.07.17.739174

**Authors:** Vanessa Klöker, James S. Nagai, Zihao Feng, Lampros Mavrommatis, Larissa Hermanns, Jacqueline M. J. Moscoso, Mayra Ruiz, Christoph Kuppe, Ivan G. Costa

## Abstract

Single-cell sequencing has advanced the study of cell-cell communication, yet most methods focus on intercellular ligand-receptor interactions while neglecting downstream intracellular signalling cascades and the possibility that downstream target genes themselves encode ligands, thereby propagating communication across multiple cells. We present IntraTalker+CrossTalkeR that combines intracellular (IntraTalker) and intercellular (CrossTalkeR) signalling from multimodal single-cell data. IntraTalker infers cell-type-specific transcription factor activities and constructs receptomes that link receptors to downstream target genes, which are then integrated with ligand-receptor predictions in CrossTalkeR. To prioritize signalling receptors, the framework performs in silico receptor perturbation. In a murine bone marrow dataset, this recovered the known function of the Il1r1 receptor in driving myeloid progenitor cell-state changes. In a human myocardial infarction dataset, it predicted a novel role for IGF1R signalling in driving the differentiation of fibroblasts towards progenitor fibroblast states.

## Introduction

Recent advances in single-cell sequencing methods, such as single-cell RNA sequencing (scRNA-seq) and single-cell ATAC sequencing (scATAC-seq), enable a detailed understanding of complex biological processes, including cell-cell communication, transcriptional regulation, and cell differentiation^1–3^. The great majority of cell-cell communication methods explore intercellular communication through paracrine signalling, specifically ligand and receptor interactions among cell types detected in single-cell data sets ^4,5^. However, complex diseases and tissue processes, such as fibrotic tissue remodelling^6^, are consequences of a cascade of intercellular paracrine signalling that leads to a cellular response via intracellular signal transduction. Moreover, the latter can further generate paracrine signalling affecting surrounding cells. Despite the importance of considering both intracellular and intercellular signals, very few approaches address this problem holistically and integratively^7^.

A common approach to introduce intracellular signalling is to leverage pathway databases to relate receptors and transcription factors (TF) and to estimate receptor or pathway activity by considering the TF activity of downstream transcription factors, as explored in scMLnet^8^, scSeqComm^9^, CellphoneDB v.4^10^, and LIANA+^11^. TF activity is typically defined as a statistic that quantifies the concordance and strength of the expression patterns among a TF’s target genes, where TF-target gene interactions are organized into regulons, usually derived from a priori gene regulatory network databases such as Reg Network or TRRUST ^12, 13^. A key advantage of regulon-based TF activity is its footprinting property. In the absence of direct measurements of protein abundance that characterize pathway activation, transcriptional patterns of TF target genes can serve as a proxy for measuring the activity of a TF and upstream signalling components^14,15^. Despite sharing similar concepts, scMLnet, scSeqComm, Liana+, and CellPhoneDB differ substantially in the statistical methods used to measure TF activity, the regulon databases they employ, and their conceptual focus. While all these distinct approaches capture the relationships between intercellular communication and intracellular signalling, they do not account for the fact that target genes may encode ligands, thereby enabling additional intercellular communication.

Multimodal sequencing and integrative analysis of transcriptional (scRNA-seq) and chromatin structure (scATAC-seq) enable the generation of cell-type-specific gene regulatory networks (GRN)^16,17^. Nevertheless, all existing intracellular communication approaches rely on cell-agnostic TF regulons, thereby ignoring cell-type-specific chromatin structure, which is crucial in gene regulation processes^18^. Besides enabling the analysis of regulatory mechanisms controlling cell differentiation^19,20^, GRNs can also be used to simulate gene expression perturbation experiments. In this context, machine learning models are trained using upstream TF activity as a feature to predict target gene expression. These models can then be used to investigate whether perturbations such as knock-outs (KO) or stimulation of a TF in these networks induce expression changes that drive cell state transitions (CellOracle^21^ and SCENIC+^20^).

While previous approaches couple ligand-receptor inference with transcription factor activity scoring (Supp. table S1), they do not construct explicit receptor-to-target-gene signalling paths, and therefore do not capture the possibility that downstream target genes themselves encode ligands that propagate communication to further cells. To address this gap, we propose the IntraTalker+CrossTalkeR2 framework, which builds upon the graph-based analysis of intercellular communication (CrossTalkeR^22^) by integrating intracellular signalling graphs to improve the prioritization of cells, ligands, receptors, and TFs driving cell-cell communication. IntraTalker uses single-cell transcription factor activities, either inferred from prior knowledge or derived from multiome-derived gene regulatory networks. Moreover, IntraTalker introduces the concept of receptomes, linking receptors to downstream target genes through transcription factors and enabling modelling of receptor perturbation simulations and transitional modelling of cell states. We applied the IntraTalker+CrossTalkeR2 framework to two distinct biological case studies. The first is based on a public available murine bone marrow ligand knockout scRNA-seq data set investigating the role of the endogenous interleukin-1 receptor antagonist (Il1rn) in regulating interleukin-1 beta (Il1b) signalling^23^. The second case study was performed on a publicly available human multi-omic heart data set focusing on cardiac remodelling upon myocardial infarction^24^, where we could recapitulate the known role of TGFB1 in fibrosis and predict a potential novel role of IGF1R signalling in driving fibroblasts towards progenitor states.

## Results

### Overview of IntraTalker+CrossTalkeR2

IntraTalker+CrossTalkeR2 is a framework for integrating inter- and intracellular communication by leveraging multimodal single-cell data (scRNA-seq and scATAC-seq) and performing in silico receptor perturbation simulations (Fig. 1). The framework consists of three main modules: (1) analysis of intracellular signalling; (2) integration with intercellular communication; (3) simulation of receptor knockout (KO) or over-expression based on inferred cellular signalling. First, we identify condition- and cell-type-specific transcription factor activities. This is achieved using a prior-knowledge regulon database^25^, or, if available, gene regulatory networks (GRNs) inferred from scATAC-seq data^16^. Next, pathway and ligand-receptor interaction databases^26^ are used to link transcription factors to upstream receptors and to transcription factor target genes, thereby constructing intracellular signalling networks. These networks are denoted as receptomes, as they connect receptors to downstream target genes. Ligand-receptor interaction predictions^11^ and intracellular networks from IntraTalker are then analyzed with CrossTalkeR2. This allows us to identify key ligands, receptors, transcription factors, and cells involved in cell-cell communication signalling in disease vs. normal conditions. Notably, CrossTalkeR2 introduces novel features, such as the identification of cell communication communities by applying community detection algorithms^27^ to directed, weighted, and signed cell-cell communication networks^28^. The third main module of the IntraTalker+Cross TalkeR2 framework focuses on in silico receptor perturbation experiments. We leverage the receptomes to train machine learning models that predict downstream gene expression from the receptor expression activity, similar to transcription factor in silico perturbation approaches^20^. These machine learning models are used to simulate changes in expression under receptor perturbations (stimulation or KO). The results are projected onto cellular trajectories, which allow us to rank receptors regarding their ability to change cell states.

**Figure 1.**
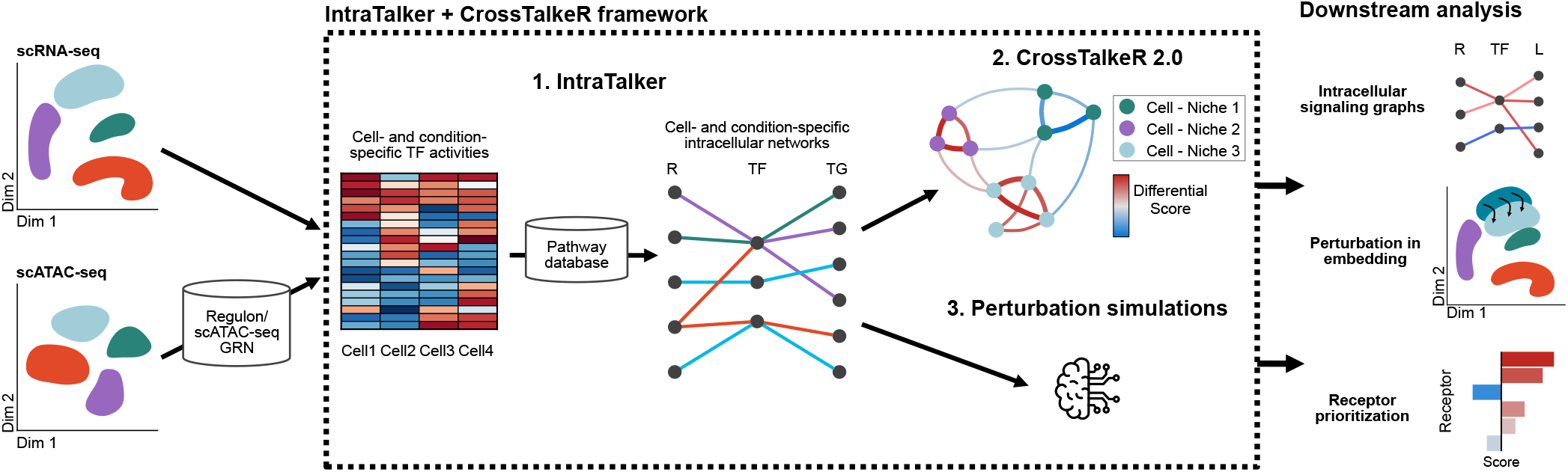
Overview of the IntraTalker+CrossTalkeR2 framework. Multimodal single-cell data (scRNA-seq and scATAC-seq) is given as input to IntraTalker+CrossTalkeR2 framework. In the first module, it infers cell-type-specific regulatory activity and constructs intracellular signalling networks. Next, these are integrated with ligand-receptor predictions to model cell-cell communication and identify key signalling components and cellular communities with CrossTalkeR2. Finally, the in silico receptor perturbation module predicts the effects of receptor signal knockouts or stimulations on cellular states. The framework returns intracellular signals and projections reflecting the effects of perturbations and ranks receptors by their ability to drive cell-state changes.

### Effects of Il1rn Knockout in Bone Marrow Cell-Cell Communication

To evaluate the power of IntraTalker+CrossTalkeR2 for characterizing cell-cell communication events and to rank receptors, we re-analyzed a publicly available scRNA-seq dataset with myeloid, stromal, and progenitor populations from mouse bone marrow collected under Il1rn knockout (KO) and wild-type (WT) conditions. There, Villatoro et al.^23^ showed that the reduced activity of the Il1r1 antagonist ligand Il1rn is linked to increased Il1b signalling, resulting in altered hematopoietic progenitor cell-states, increased myeloid expansion, and Il1b-induced damage to mesenchymal stromal cells (MSC) (Sup. Fig. S1a). Patient studies have shown that this mechanism is associated with myeloproliferative diseases and acute myeloid leukemia (AML)^29^.

We integrated the data, while preserving the original cell types annotation, including twenty cell types (Fig. 2a). We next examined the expression of interleukin genes and observed condition-dependent expression between the WT and KO conditions for the ligand *Il1b*, the antagonist ligand *Il1rn*, and the receptor *Il1r1* (Sup. Fig. S1b). In particular, we observe a decrease in Il1rn expression in MSCs and G4 neutrophils, along with an increase in the expression of the antagonized receptor Il1r1 in MSCs, confirming the effect of the KO experiments.

**Figure 2.**
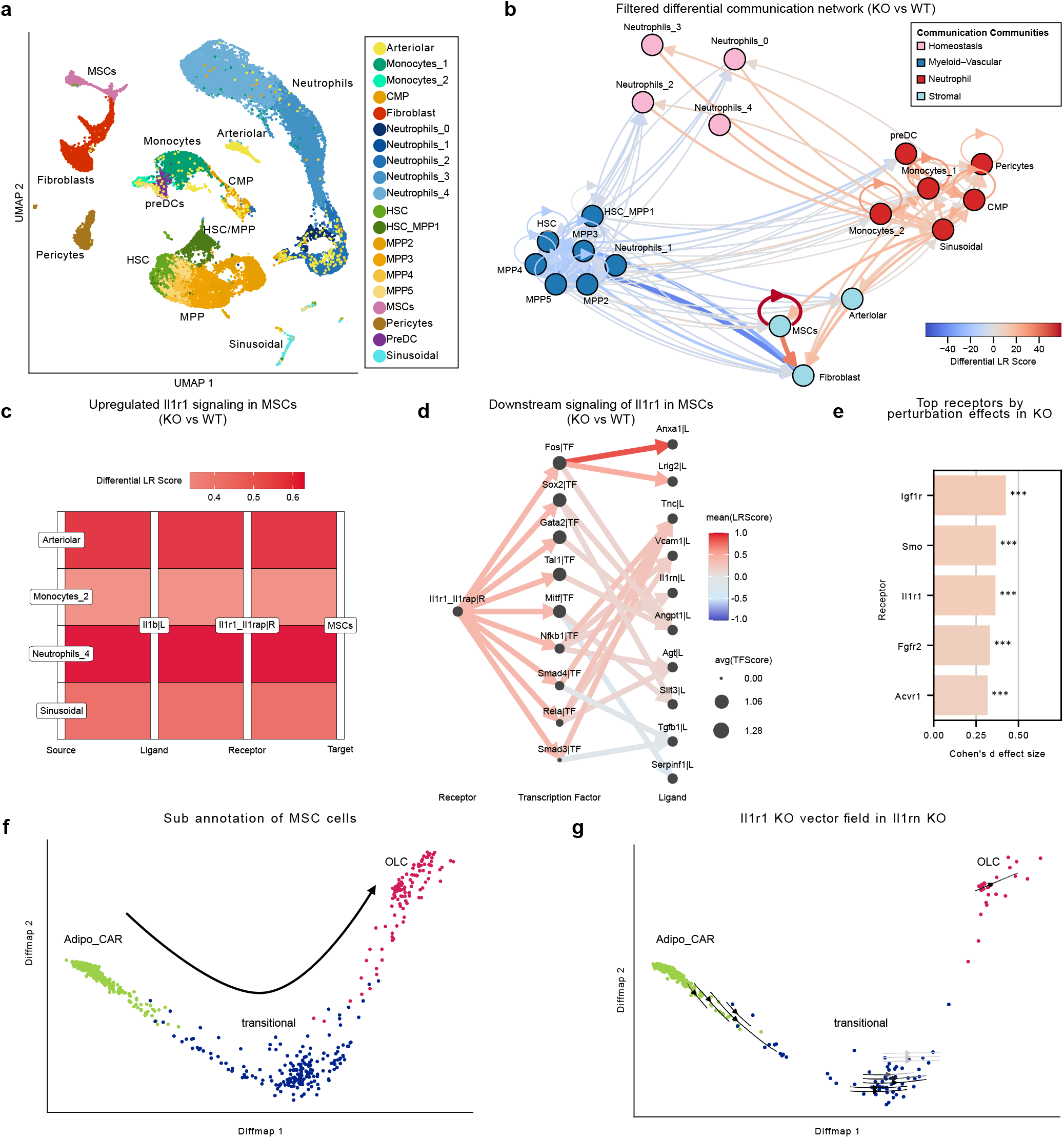
**a** UMAP of the integrated single-cell transcriptome data with 20 cell types (n=35,435). MSCs, mesenchymal stem cells. pre-DCs, precursor dendritic cells. CMP, common myeloid progenitors. MPP, multipotent progenitors. HSC, hematopoietic stem cells. **b** Differential cell-cell interaction plot visualizing identified communication communities in Il1rn KO versus WT conditions. The node color indicates the affiliation with a communication community. The edge width and color correspond to the LR interaction intensity. Edges were filtered by Fishers exact test on cell proportions with *P*-value < 0.05. **c** Sankey plot showing all detected Il1b - Il1r1 iterations in KO condition. **d** Intracellular signalling graph highlighting downstream signalling of Il1r1 in MSCs. The TF node size reflects the average TF activity score. The color scale indicates the relative mean LR score of the incoming receptor and outgoing ligands. **e** Top five receptors by perturbation score (Cohens’ d) considering all MSC cells. * * * *P* < 0.001 (Wilcoxon signed-rank test). **f** Diffusion map representation of MSCs sub-clusters (AdipoCAR, transitional and OLCs, n=828). The arrow indicates the trajectory from AdipoCAR towards OLCs. OLC, osteolineage cells. **g** Predicted shifts after Il1r1 KO simulation in MSC cells in the Il1rn KO condition. Arrows are shaded by the distance each cells shifted after receptor simulation.

We next used Intra+CrossTalkeR2 to characterize cell-cell communication by comparing KO and WT conditions. This delineated four cell-cell communication communities: one with a decrease in communication in the KO condition, mostly containing progenitor cells (stem/progenitor community), and three communities with an increase in communication: one containing myeloid cells, one with neutrophils, and one with stromal cells (Fig. 2b). In particular, MSCs are the cell type with the highest signals within the bone marrow network upon Il1rn KO (Sup.Fig. S1c). Interactions between stromal cells are predominantly upregulated, whereas there is only a slight upregulation from progenitor cells towards MSCs (Sup. Fig. S1c). Especially incoming interactions from arteriolar and sinusoidal cells are of interest in accordance with the mechanisms proposed in^23^. If we focus on interleukin-related Il1r1 signalling, it is predicted to be exclusive toward MSCs (Fig. 2c). As expected, Il1b – Il1r1 interactions are predicted to be upregulated following the KO of Il1rn, driven by incoming signals from arteriolar endothelial cells and G4 neutrophils. Assessing downstream signalling of Il1r1 (Fig. 2d) reveals the transcription factor Nfkb1 as a mediator of Il1rn activation, implicating the involvement of the Nfkb-pathway, as already supported ^23^. Further intracellular signalling includes fibrosis-related SMAD3/4 transcription factors, which are predicted to up-regulate the fibrosis-related TNC gene^30^. Altogether, these results indicate that IntraTalker+CrossTalkeR2 can delineate relevant cellular communities, ligand-receptor pairs, and downstream signalling cues associated with interleukin signalling.

We next evaluated the in silico perturbation models to predict receptors associated with changes in MSC disease states. Clustering of MSC cells identified three main sub-clusters (Lepr+ adipogenic CAR cells (Adipo-CAR), osteolineage-committed (OLC) cells, and transitional-state cells), which were characterized by their gene markers and label transfer from a bone marrow atlas^31^(Sup.Fig. S1d). The trajectory and pseudotime analysis revealed an axis from Adipo-CAR cells via transitional cells to OLCs (Fig. 2f; Sup.Fig. S1e). We performed IntraTalker-guided receptor KO analysis with 30 receptors predicted to be active in MSCs and associated with downstream regulomes. Receptors were then ranked based on their capacities to induce cell differentiation shifts, quantified as an increase or decrease in pseudotime upon perturbation (perturbation score). Here, a positive receptor perturbation score indicates a cell state change towards OLCs, while a negative score indicates a change towards Adipo-CAR cells.

Given our focus on fibrosis-associated OLC differentiation, receptors with the largest positive changes are most relevant here. Notably, Il1r1 ranks third regarding the change in perturbation score (Cohen’s *d* effect size; Fig. 2e). As expected, the effect of Il1r1 perturbation simulation is more pronounced in cells measured under Il1rn KO condition than in cells in the Il1rn WT condition (Sup. Fig. S1f-g), reflecting the activation of Il1r1 upon the KO of its antagonist Il1rn. To visualize the effects, perturbation scores were projected as vector fields along the trajectory, following a previous approach^20^. This particularly highlights an increase in the cell-differentiation towards transitional and OLCs cells upon Il1rn KO (Fig. 2g). Altogether, these results show that IntraTalker receptor perturbation analysis identifies Il1r1 receptor as an important signalling driver in the transition of Adipo CAR cells towards OLCs.

### Signalling Relevant to Fibroblast Differentiation in Myocardial Infarction

The spatial atlas of myocardial infarction^24^ provides a high-resolution multiomic map including snRNA-seq and snATAC-seq of cardiac remodelling after myocardial infarction, showing how fibrosis impacts remodelling of the heart tissue. We analyzed the publicly available snRNA-seq and snATAC-seq data by considering myogenic (control) and ischemic samples from the study using our IntraTalker+CrossTalkeR2 framework to characterize the main cellular and cell-cell communication drivers of fibrosis. Here, we leveraged the snATAC-seq data to generate cell-specific eGRNs, which were estimated with scMega^16^ to be used as input to IntraTalker+CrossTalkeR2.

The differential cell-cell communication analysis comparing the ischemic and myogenic conditions detected six communication communities (Fig. 3a), of which only one community was associated with increased signalling in the ischemic condition. This included lymphatic endothelial cells, mast cells, myofibroblasts (Myofib), C7^+^ fibroblasts (C7^+^Fib), and damaged cardiomyocytes. This is in line with previous results from Kuppe et al. ^24^, where high signalling activity between myofibroblasts and damaged cardiomyocytes was observed.

**Figure 3.**
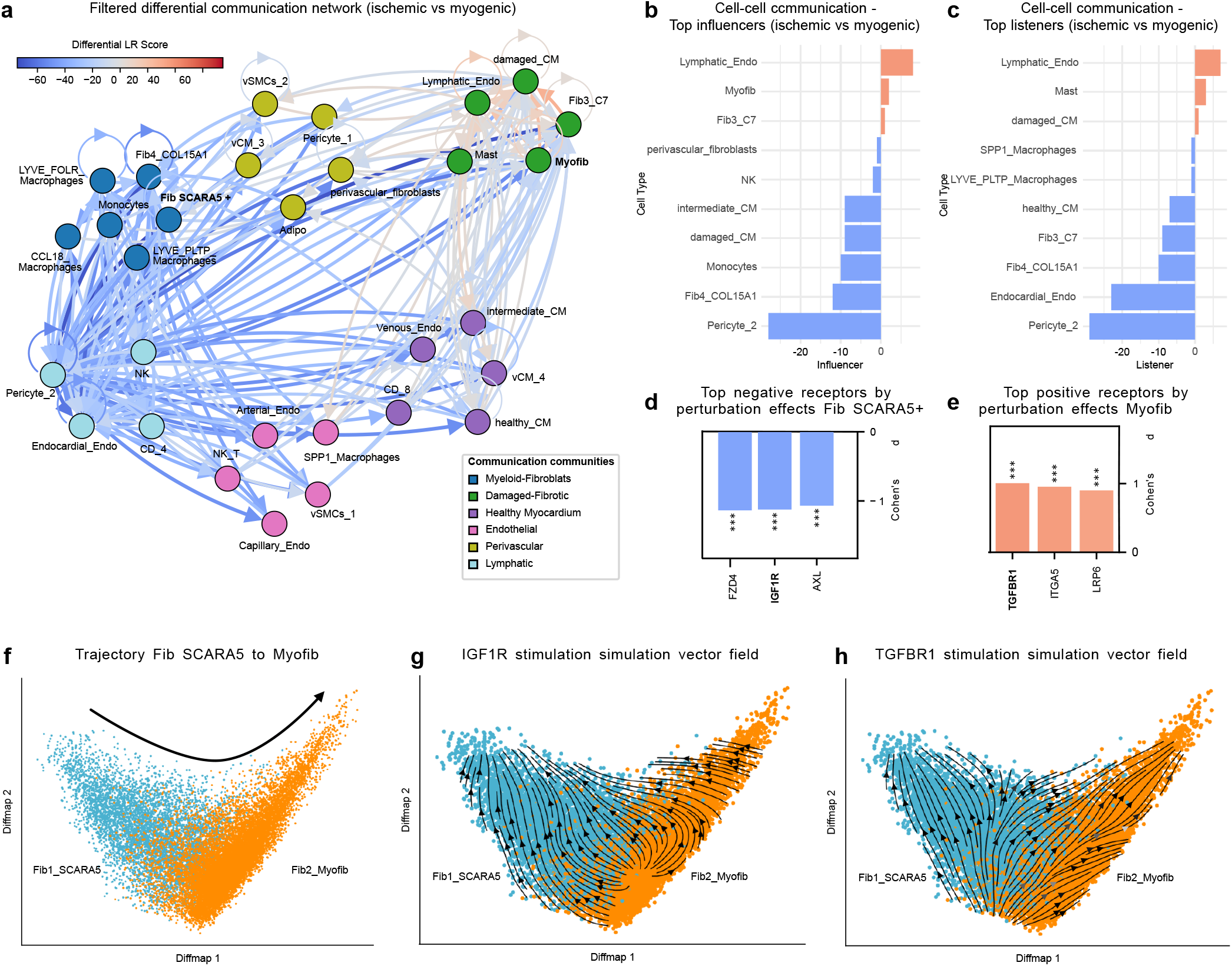
**a** Filtered differential cell-cell interaction plot visualizing identified communication communities in ischemic versus myogenic conditions. The node color indicates the affiliation with a communication community. The edge width and color correspond to the LR interaction intensity. Edges were filtered by Fishers exact test on cell proportions with a *P*-value cutoff of 0.05. **b** Top ten influencer scores in the cell-cell communication network comparing ischemic against myogenic conditions. **c** Top ten listener scores in the cell-cell communication network comparing ischemic against myogenic conditions. **d** Top 3 receptors by negative perturbation score (Cohens’ d) considering shifts in SCARA5^+^ fibroblasts. * * * *P* < 0.001 (Wilcoxon signed-rank test). **e** Top 3 receptors by positive perturbation score (Cohens’ d) considering shifts in the myofibroblasts. * * * *P* < 0.001 (Wilcoxon signed-rank test). **f** Difussion map representation of SCARA5^+^ fibroblasts and myofibroblasts. The arrow indicates the trajectory from SCARA5^+^ fibroblasts towards myofibroblasts. **g-h** Predicted shifts after FZD4 (g) and TGFBR1 (h) stimulation in SCARA5^+^ fibroblasts and myofibroblasts. Arrows are shaded by the distance each cells shifted after receptor simulation.

Even though positive signalling associated with the ischemic condition was sparse, network ranking analysis from CrossTalkeR2 revealed lymphatic endothelial cells, myofibroblasts and C7^+^ fibroblasts as top influencers — cells with high out going signals — in the ischemic condition (Fig. 3b-c). Among these populations, the prominent roles of myofibroblasts and lymphatic endothelial cells are in line with their associated mechanisms in cardiac remodelling. Myofibroblasts are the primary source of extracellular matrix proteins that stabilize the myocardium following cardiomyocyte death^32,33^, whereas lymphatic endothelial cells are key cells in lymphangiogenesis, and are known to invade the damaged tissue areas^34^. Both mechanisms can be activated by mast cells^35^, which were also associated with the ischemic condition, with the release of major fibroblast growth factors and lymphangiogenesis activating signalling.

To dissect the mechanisms driving fibrosis, we focused on signalling pathways in fibroblasts. In particular, we seek to understand how intercellular signals and their downstream intracellular pathways promote the differentiation of progenitor fibroblasts, such as SCARA5^+^ fibroblasts^24^, into a myofibroblast phenotype (Fig. 3f;Sup.Fig. S2a). We performed trajectory analysis of SCARA5^+^ fibroblasts and myofibroblasts, as well as an in silico perturbation analysis of 27 receptors associated with cell–cell communication targeting these fibroblast populations. Considering the receptors with the three highest positive perturbation scores, corresponding to increased differentiation towards myofibroblast state, and the three lowest scores, corresponding to transitions toward the SCARA5^+^ fibroblast state, several biologically relevant receptors were identified (Fig. 3d-e). Among the receptors predicted to promote myofibroblast differentiation is TGFBR1, a key mediator of TGF-*β* signalling^36^ (Fig. 3h). In contrast, receptors predicted to favor transitions toward the SCARA5^+^ state included the WNT receptor FZD4 and IGF1R(Fig. 3g), both of which are associated with fibrosis-related signalling pathways^37^. Of note, LRP6, which is a repressor of canonical WNT signalling^38^, is predicted to drive fibroblast towards myofibroblast states.

To investigate cell-cell communication mediated by the receptors of interest, we examined incoming intercellular signalling directed toward SCARA5^+^ fibroblasts and myofibroblasts within the myeloid-fibroblast and damaged-fibrotic communities, respectively, revealing signalling patterns characteristic of each population (Sup.Fig. S3a-b). SCARA5^+^ fibroblasts received signalling associated with the myogenic condition. This includes IGF1-IGF1R predominantly by LYVE1^+^ resident macrophage subsets, in line with this macrophage population regulating tissue homeostasis^39,40^; active WNT2B-FZD4 WNT signalling, where WNT2B might be involved with non-canonical WNT signalling^41^ (Sup.Fig. S3a). In contrast, myofibroblasts showed enhanced TGFB1-TGFBR1 signalling, one main driver of fibroblast-to-myofibroblast differentiation^42^ (Sup.Fig. S3b).

Enrichment analysis of the downstream ligands (Sup.Fig. S3c-d) with GO Biological Process ^43,44^ further supported these cell-state-specific signatures (Sup.Fig. S3e-f). Down-regulated SCARA5^+^ fibroblast ligands were dominated by vascular and barrier-supportive programs, including vasculature development (ANGPT1, THBS2, VEGFA, CXCL12, SEMA3C, THBS1, JAG1) and regulation of endothelial cell differentiation (VEGFA, VCL, JAG1). In contrast, up-regulated myofibroblast ligands defined an active wound-healing and remodelling program. The top terms were positive regulation of cell migration and response to wounding (including HGF, VEGFC, CCN4, TNC, SPP1, THBS1, TGFB1) together with extracellular matrix organization (POSTN, LAMB1, COL12A1, COL16A1, COL27A1, TGFB1), matching the migratory, matrix-depositing phenotype that builds the fibrotic scar^42^. Notably, myofibroblast ligands were additionally enriched for osteoblast differentiation and ossification (SPP1, CCN4, TNC, IGF1, CCN1), echoing the bone- and cartilage-like ECM program of matrifibrocytes, which were described as highly differentiated fibroblasts^45^. These results suggest distinct signalling by these two fibroblast cell states.

### Validation Experiments on Human Heart-Derived Fibroblasts

To validate the previous predictions, we cultured human heart-derived fibroblasts and treated them with two ligands associated with two predicted receptors: TGFB1 as a ligand related to the stimulation of the myofibroblast state via TGFBR1 and IGF1 as a ligand associated to drive fibroblasts towards the SCARA5^+^ fibroblast states via signalling via IGF1R. First, we selected gene markers associated to these two fibroblast sates^46^ (Supp. Table. S2). Regarding SCARA5^+^ fibroblasts, we select the markers PCOLCE2 (procollagen C-proteinase enhancer 2), which is a strong, cross-tissue marker of quiescent mesenchymal/stromal cells^47^, EBF1, which is a transcription factor related to perivascular cells^48^; and GRK5 (G protein-coupled receptor kinase 5 ), which expression rescue has been shown ameliorate fibrosis^49^. For myofibroblasts, we select periostin (POSTN) and fibronectin (FN1) as two as pan-fibrosis myofibroblast/ECM markers.

We then used qPCR to validate the expression changes of these genes. TGFB1 stimulation increased the expression of POSTN and FN1, while decreased the expression of EBF1, PCOLCE2 and GRK5 ( Fig. 4) supporting the well-known role of TGFB1 in fibrosis and the selected markers. Regarding IGF1, we observe that while it increased the expression of all markers, we observe a particular increase in SCARA5 fibroblast markers as PCOLCE2 and GRK5. These results indicates that IGF1 stimulation is able to trigger the expression program of SCARA5^+^ fibroblast cells, while the expression of this program is reduced by the TGFB1, as predicted by CrossTalkeR2+IntraTalker.

**Figure 4.**
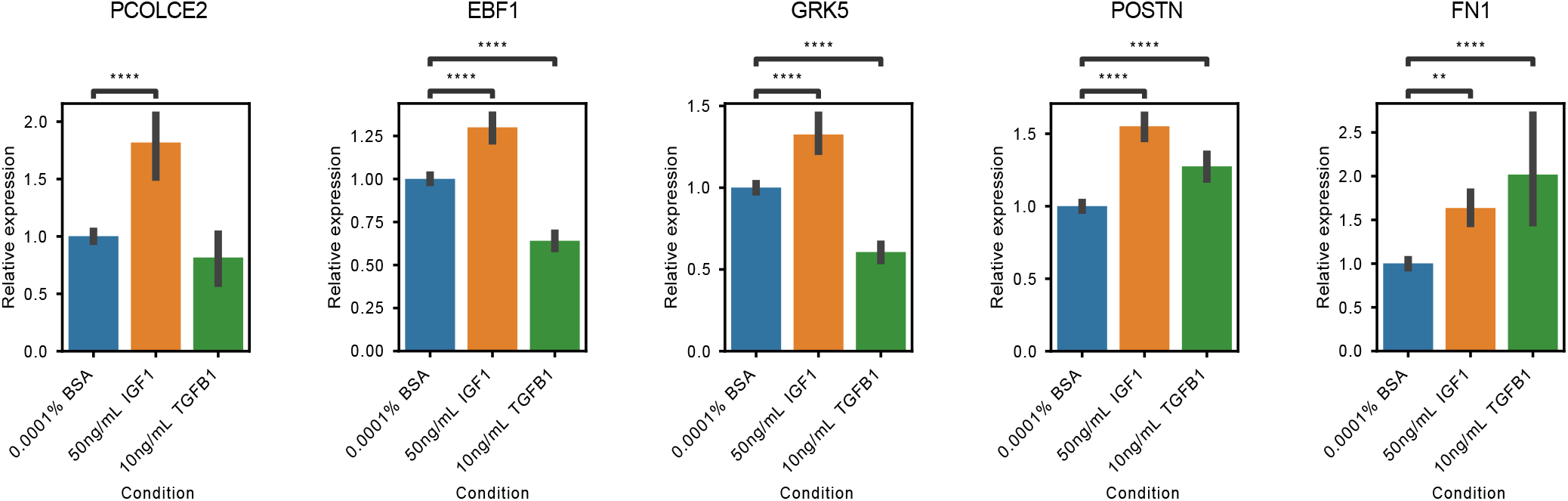
qPCR experiment with the expression of SCARA5^+^ Fib. and Myofibroblasts stimulated with IGF1 and TGFB1 on three biological and three technical replicates. Differences in relative gene expression (using BSA as reference) were compared using a linear mixed model; adjusted p-values are indicated as follows: ^*^ *p* < 0.05, ^**^ *p* < 0.01, ^***^ *p* < 0.001 and, ^****^ *p* < 0.0001.

## Discussion

Cell-cell communication is driven not only by ligand-receptor interactions between cells but also by intracellular downstream signalling that mediates cellular responses and propagates communication across cells. While several existing frameworks already couple ligand-receptor inference with transcription factor activity scoring, IntraTalker+CrossTalkeR2 extends this integration in three ways: by constructing explicit receptor-to-target-gene receptomes that allow downstream target genes to re-enter the network as ligands for further intercellular signalling; by incorporating cell-type-specific gene regulatory networks derived from scATAC-seq data rather than relying solely on a priori regulomes; and by coupling this intracellular signalling layer to in silico receptor perturbation simulations that quantify receptor impact on cell-state transitions along a trajectory. The resulting signalling graphs can be used to prioritize cells, rank receptors and detect cellular communities. Moreover, in silico receptor perturbations allow the further delineation of actionable targets for potential therapeutics.

Analysis of the Il1rn KO mouse bone marrow study highlighted MSCs as the central receivers of interleukin signalling via Il1b. The framework recovered canonical NFkB downstream signalling as well as fibrosis-associated intracellular signalling involving SMAD3/4 and the TNC ligand. Moreover, in silico perturbation prioritized Il1r1 as one of the top functional drivers and predicted an MSC state transition towards the osteo-lineage, which is in agreement with pathological remodelling in the fibrosis context^50^.

The application in the myocardial infarction study identified a cellular communities associated with cardiac remodelling, including myofibroblasts, lymphatic endothelial cells, damaged cardiomyocytes, and mast cells, which closely reflect mechanisms of fibrotic remodelling in the heart^24^. Moreover, in-silco perturbation captures bona fide fibrosis-related receptors, such as TGFBR1. More importantly, we used an in-vitro fibroblast cell culture to observe the role of a novel putative signalling pathway, IGF1R, which was predicted by IntraTalker. Our finding that IGF1R signalling promotes a SCARA5+ progenitor-like fibroblast state is consistent with prior work showing that myofibroblast-specific deletion of Igf1r worsens interstitial fibrosis in a mouse model of angiotensin II/phenylephrine-induced cardiac fibrosis, whereas low-dose IGF-1 infusion attenuates fibrosis by suppressing fibroblast proliferation and differentiation into aSMA+ myofibroblasts^51^. Notably, the role of IGF1R in cardiac fibrosis appears to be context- and cell-type-dependent: while cell-type-specific loss of IGF1R in fibroblasts exacerbates fibrosis, global heterozygous Igf1r deficiency has instead been reported to attenuate angiotensin II-induced fibrosis via downregulation of the Akt/ERK/NFKB pathway, in which IGF1R directly interacts with GRK5^52^. This latter study identified GRK5 as a direct interacting partner required for IGF1R’s downstream signalling in cardiac remodelling, which parallels our own observation that IGF1 stimulation increased GRK5 expression alongside other SCARA5+ markers in cultured human cardiac fibroblasts, and suggests IGF1R-GRK5 signalling as a specific axis worth further mechanistic dissection.

While IntraTalker+CrossTalkeR2 offers an integrative framework for intra- and intercellular communication, some limitations warrant future development. Inferred signalling relationships remain indirect, lacking to fully capture pathway complexity. Integrating protein-level measurements and pathway activity inference could improve receptor activity modelling, but the lack of single-cell protein measurements does not allow such studies. As for the perturbation simulations, the current framework does not explicitly model signalling kinetics or nonlinear regulatory interactions. Further benchmarking of cell-state-transition-based methods remains a challenge to be tackled in the future.

## Methods

### CrossTalkeR2 + IntraTalker

IntraTalker+CrossTalkeR2 is a framework for combining inter- and intracellular communication by considering multi-modal single-cell data (scRNA-seq and scATAC-seq) and performing in silico receptor perturbation simulations. We will describe below the three main methodological parts of the framework, which include the intracellular communication (IntraTalker), the intercellular communication (CrossTalkeR2), and the in silico receptor perturbation approach.

### IntraTalker

The framework requires preprocessed and normalized scRNA-seq data, along with cell cluster and condition information. Let *X* ∈ ℝ^*C*×*G*^ denote the single-cell matrix, where *C* is the number of cells and *G* the number of genes. Let *K* be the total number of cell clusters. A cell belonging to a cell cluster can be represented as *c*_*i*_ ∈{1, …, *K* }.

IntraTalker first constructs an intracellular signalling network to relate receptors to downstream transcription factors, based on prior knowledge. We utilize the Omnipath meta-database, which integrates multiple pathway resources, including curated databases, experimental studies, and computational predictions^26^. We restrict the protein-protein interaction network to retain only intracellular signals by excluding upstream ligand-receptor interactions and all interactions downstream of transcription factors. As a result, we will obtain a directed graph linking all available receptors and transcription factors in Omnipath. Receptors have a directed edge towards a TF, whenever we detect a path between the receptor and the TF.

For transcription factor analysis, IntraTalker relies on regulomes (transcription factor-target gene interactions), provided either from the cell-agnostic database DoRothEA^25^ or, optionally, cell-type-specific regulomes can be inferred from multimodal (RNA+ATAC) single-cell data using scMEGA^16^. A cell-agnostic regulome is represented as a list of triplets *R*_*Regulome*_ = {*r*_1_, …, *r*_*n*_}, where each triplet *r* = (*source, target, weight*) consists of a transcription factor *source*, a target gene *target*, and a interactions *weight* ∈ [−1, 1], indicating the strength and mode of action (activator or repressor). To extend the regulome, we integrate cell-type-specific gene regulatory networks derived from scATAC-seq data, represented as *R*_*GRN*_ = {*r*_*n*+1_, …, *r*_*n*+*m*_ }. The final regulome is then defined as:

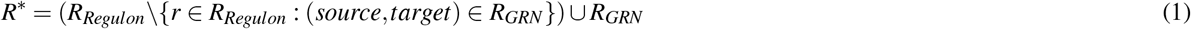

Thus, interactions present in both sets are replaced by those from *R*_*GRN*_, while novel interactions from *R*_*GRN*_ are added. This approach emphasizes transcription factor–target gene interactions inferred from scATAC-seq data over prior-knowledge regulomes. Of note, DoRothEA provides interactions annotated with curation levels A–E, each level reflecting the supporting evidence. Here, we use the most stringent interactions of levels A and B, which are supported by curated resources or a supported multiple lines of experimental evidence, as the default.

Intra Talker performs transcription factor activity predictions using decoupleR^53^, which estimates transcription factor activity from the scRNA-seq expression matrix *X* based on weighted regulomes. It uses the univariate linear model (ULM) as recommended, obtaining transcription factor activity scores at single-cell resolution suitable for downstream analysis with Seurat or anndata objects. For datasets with multiple conditions, it uses cluster-wise pairwise *t*-tests with the scran package and computed r effect sizes to assess differential transcription factor activities. For single-condition datasets, IntraTalker uses Seurat’s FindAllMarkers function with the Wilcoxon test to compare each cell type against all others. In both cases, the default significance filter for transcription factors is the thresholds *p* < 0.05 (Benjamini-Hochberg adjusted) and |*logFC*| > 1.

Finally, cell-type-specific active transcription factors are linked to upstream receptors and downstream target genes using the prior knowledge resources. This results in a set of 4-tuples *Y* ={ *y*_1_, …, *y*_*n*_ }, we refer to as receptome. Each element is defined as

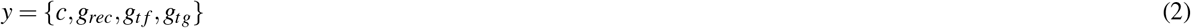

where *c* ∈ *K* denotes the cell type, *g*_*rec*_ a receptor gene, *g*_*t f*_ a transcription factor gene, and *g*_*tg*_ a target gene. Target genes are retained only if expressed (exp(TG) > 0). The receptome is the core structure of the framework, supporting downstream signalling analysis and receptor perturbation simulations.

### CrossTalkeR2

To integrate intracellular signalling with intercellular communication, we combine the inferred receptome with predicted ligand-receptor interactions. Ligand-receptor interactions were obtained using the LIANA+ framework^11^ and represent the intercellular signalling layer connecting sender and receiver cell types. Following CrossTalkeR^22^, LR interactions are defined as a set of 5-tuples 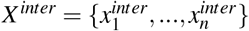, provided separately for each condition. Each tuple is defined as *x*^*inter*^ = (*c*_*s*_, *g*_*lig*_, *c*_*r*_, *g*_*rec*_, *w*), where *c*_*s*_, *c*_*r*_ ∈ *K* denote the sender and receiver cell types, *g*_*lig*_ is a ligand gene, *g*_*rec*_ is a receptor gene, and *w* > 0 the weight indicating the strength of the ligand-receptor interaction.

IntraTalker+CrossTalkeR2 integrates inter- and intracellular interactions by constraining the receptome to receptors involved in active ligand-receptor interactions. This ensures that downstream transcriptional signalling is linked to upstream intercellular communication. In addition, target genes are restricted to those associated with ligands present in the ligand-receptor interaction set, yielding a signalling structure that is consistent across both layers. We define the set of intracellular interactions as *X*^*intra*^ = *X*^*r*→*t f*^ ∪ *X*^*t f* →*l*^, where *X*^*r*→*t f*^ and *X*^*t f* →*l*^ denote the interaction from receptors to transcription factors and from transcription factors to ligands. Specifically, we define two types of 5-tuples as

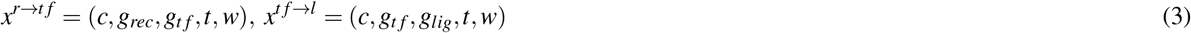

where *c* ∈ *K* is the cell type, *g*_*rec*_ is a receptor gene, *g*_*t f*_ is a transcription factor gene, *g*_*lig*_ is a ligand gene, *t*∈ {*R*→*TF, TF* → *L*}is the type of interaction, and *w* > 0 is the weight and indicates the transcription factor activity. These unified graphs, provided by *X*^*inter*^ and *X*^*intra*^, enable the propagation of signalling information across different cells, linking ligand-receptor interactions to downstream transcription factor activity. As a result, we can apply CrossTalkeR2 to perform single-condition and differential cell-cell communication analysis while accounting for intracellular signalling cascades.

Another novel aspect of CrossTalkeR2 is a community-detection approach for identifying cell-cell communication communities. Here, we consider the cell-cell networks as defined in CrossTalkeR. In short, the ligand-receptor interactions in *X*^*inter*^ are aggregated into a network with nodes representing cell types and directed edges that summarize all LR interactions between sender-receiver cell pairs. Edge weights correspond to the sum of the individual ligand–receptor interaction weights^22^. We use a kernel-similarity-based approach, which considers both directions and weights of the edges in the cell-cell communication networks ^54^, allowing the subsequent use of the Leiden community detection algorithm ^27^. Let *A* ∈ ℝ^*K*×*K*^ denote the directed network adjacency matrix, where *A*_*i j*_ represents the weighted interactions from sender cell *i* to receiver cell *j*, following the definition in CrossTalkeR^22^.

Let *v* = {1,…, *K*} denote the set of all nodes that represent interacting cells. We next embed *A* into a squared matrix *W* ∈ ℝ^*K*×*K*^ such that:

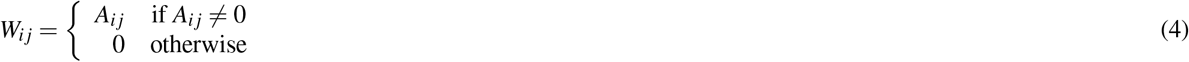

To remove scale effects associated with highly connected cell types (nodes), we applied degree normalization. The node strengths were defined as *d*_*i*_ = ∑ _*j*_ |*W*_*i j*_ |, and the corresponding diagonal degree matrix *D* = *diag*(*d*_1_, …, *d*_*K*_) was used to compute a symmetrically degree-normalized interaction matrix:

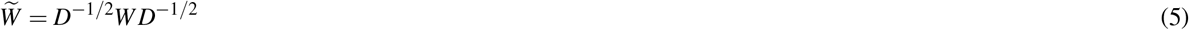

We define the kernel similarity matrix *S* based on interaction similarity as

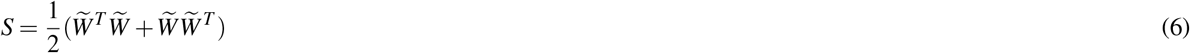

where the terms 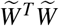 and 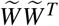 capture respectively the similarity of cells as receivers or the similarity of cells as senders.

Finally, we construct the similarity graph *G* = (*V, E, S*), where the edges *E* are defined as (*i, j*) ∈ *E* if *S*_*i j*_ > 0. Communication communities are then identified by applying community detection algorithms to the similarity graph.

For visualization, node positions are obtained using a community-informed force-directed spring layout^55^ from networkX^56^. First, communication community centers are arranged using the spring layout on the similarity graph *G*, and nodes are then positioned within each community by applying the spring layout locally. The directed edges are overlaid from the original differential network for visualization.

### Gene Expression Perturbation Module

To assess the functional impact of receptors on downstream gene expression, we implemented an in silico perturbation framework inspired by CellOracle^21^. Instead of modelling transcription factors as upstream regulators, we define regulatory relationships based on our predicted receptome, incorporating receptor-mediating signalling. Given a normalized single-cell expression matrix *X* ∈ ∈ ℝ^*C*×*G*^ and a receptome *Y*, we train gene-specific ridge regression models. First, we simplify *Y* (Eq. 2) by removing transcription factors and directly connect receptors to target genes, resulting in 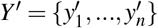 with *y*^′^ = {*c, g*_*rec*_, *g*_*tg*_}, where *c ∈ K* is the cell type, *g*_*rec*_ a receptor gene, and *g*_*tg*_ a target gene. For each target gene *g*, the expression is modeled as a function of its upstream receptors *g*_*R*_:

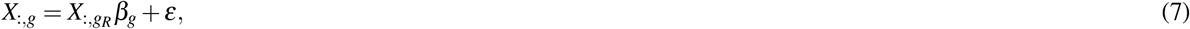

where *β*_*g*_ are the regression coefficients and *ε* is the residual error. Ridge regularization is applied with the default penalty parameter *α* = 10. The learned coefficients are aggregated into a global coefficient matrix *B* ∈ ℝ^*G*×*G*^, where each entry *B*_*i j*_ represents the influence of gene *j* in gene *i*. Non-zero entries are restricted to receptor-target gene pairs defined in the receptome.

To perform receptor perturbation simulations, the expression of the receptor of interest is modified in the initial expression matrix *X* . Knockout simulations are implemented by setting all expression values to zero, whereas overexpression is simulated by assigning a high expression value. A high value is defined as the 95th percentile of the overall expression distribution of *X* . The perturbed state is propagated through the trained regulator model by iterative matrix multiplication:

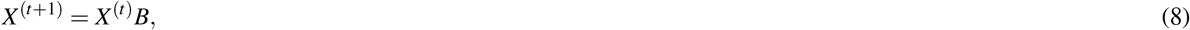

starting from the perturbed matrix *X* ^(0)^. After a defined number of iterations (*t* = 5 as default), the resulting matrix represents the simulated expression state.

To assess the effect of receptor perturbations on cellular fate, we project the perturbation effects into a low dimensional embedding space (*Z*∈ ℝ^*C*×2^) such as a diffusion map and estimate the cell pseudo-time (*t* ∈ℝ^*C*^ ) using a random walk algorithm ^57, 58^.

Next, we use a procedure from Velocyto^59^ (adapted in CellOracle^21^) to project how the expression changes observed in the perturbation experiments change the cellular state in the low-dimensional embedding. We first compute the expression difference Δ*X* = *X* ^*perturbed*^ − *X* and project the changes onto a *k*-nearest-neighbor graph constructed from the embedding coordinates *Z*. For each cell *i*, we evaluate the similarity between its difference vector Δ*x*_*i*_ and the expression differences to its neighbors (*x*_*j*_−*x*_*i*_) using a correlation-based measure. The similarity scores are converted into transition probabilities using a softmax function restricted to the local neighborhood. Finally, the displacement vector in embedding space is computed as a weighted average of normalized direction vectors towards neighboring cells, which is normalized by the local neighborhood average to remove topology-induced bias. For more details refer to CellOracle^21^.

This yields cell-wise displacement vectors Δ*Z* ∈ ℝ^*C*×*L*^, indicating the predicted directions and magnitude of the induced transitions in the low-dimensional spaces. For a continuous representation, we construct a smoothed vector field over a regular grid in the embedding space. For each grid point *z* ∈ *Z*, we compute a weighted average of the nearby displacement vectors:

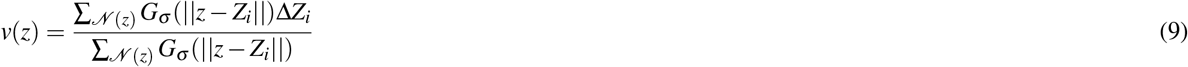

where *G*_*σ*_ is a Gaussian kernel and *N* (*z*) denotes the local neighborhood.

Moreover, we further quantify the perturbation effect using the pseudotime. We estimate the local pseudotime gradient along the displacement vectors. For each cell *i*, the gradient ∇*pt*_*i*_ ∈ ℝ^2^ is calculated by fitting a linear model over its nearest neighbors:

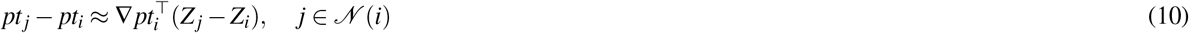

The resulting gradients are normalized to unit vectors and represent the direction of increasing pseudotime. We project the displacement vectors for each cell *i* onto the pseudotime gradients to calculate the differential pseudotime:

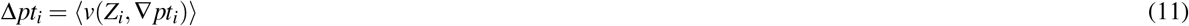

where *v*(*Z*_*i*_) denotes the vector field evaluated at the position of cell *i*. Positive differential pseudotime indicates progression along the trajectory, whereas negative values indicate opposing shifts along the trajectory. The perturbed pseudotime is then defined as

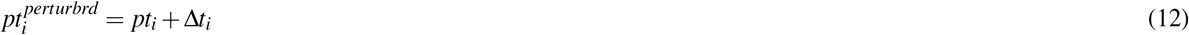

To finally quantify the perturbation effects, we compute the differences between perturbed and original pseudotime values and summarize them using the Wilcoxon signed-rank test and Cohen’s *d* effect size statistics. We evaluate the receptors based on their *p*-value significance level and the effect size, where *d* is interpreted as negligible (|*d*| < 0.2), small (0.2 ≤ |*d*| < 0.5), moderate (0.5 ≤ |*d*| < 0.8), and large (|*d*| ≥ 0.8) as standard.

### Single-Cell Integration and Communication Analysis of Il1rn Knockout Bone Marrow Data

We retrieved three Seurat v3 scRNA-seq objects corresponding to myeloid, LSK, and stromal cell experiments from the GitHub repository provided by Villatoro et al.^23^. These objects contain raw and normalized counts, along with sample annotations and cell cluster identities. First, we updated the objects to Seurat (v5.3.1) v5^60^ assays and reintegrated the samples within each dataset. We scaled the data and performed principal component analysis (PCA) and UMAP dimensionality reduction using default parameters. We integrated the provided samples for each dataset using the IntegrateLayers function in Seurat with canonical correlation analysis (CCA), following the standard Seurat workflow. To integrate the three datasets, we used Harmony^61^ with Seurat to compute a joint UMAP embedding. For downstream analysis steps in Python, we converted the joint Seurat object into AnnData^62^ format.

We inferred ligand-receptor interactions using CellphoneDB v2.0^63^ via the LIANA+ framework (v1.6.1)^11^ and the mouse consensus database^11^, with the LR expression proportion threshold set to 0.1. Predicted ligand-receptor interactions with *p* < 0.05 were considered for downstream analysis. We estimated transcription factor activity using the ULM method from decoupler (v2.1.2)^53^ and the DoRothEA regulome^25^ restricted to confidence levels A and B. We retrieved cell-type and condition-specific transcription factor activities using IntraTalker by comparing Il1rn knockout and wild-type conditions. Transcription factors were filtered by *p* < 0.01 and *LFC* > 0.4. The receptome was generated based on the retained transcription factors.

We performed differential cell-cell communication analysis comparing KO vs WT conditions using CrossTalkeR2, integrating ligand-receptor interactions with downstream transcription factor signalling. We filtered cell-cell interaction pairs using Fisher’s exact test (*p* < 0.05) and identified communication communities using the newly introduced kernel-based similarity approach, followed by Leiden (v0.11)^27^ clustering and network X (v3.5)^56^ spring layout visualization with default parameters. We performed subclustering of the MSC cluster using Scanpy’s (v1.11.5)^64^ Leiden clustering with a resolution of 0.5. We annotated cell types based on canonical lineage markers and by label transfer from the stromal bone marrow reference atlas from Baryawno et al.^31^. Label transfer was performed using Seurat’s Find Transfer Anchors and Transfer Data functions in R. We performed trajectory inference using diffusion map analysis (diffmap)^65^, followed by pseudotime estimation with diffusion pseudotime (dpt)^66^ with Scanpy. We defined the root cell population as Adipo-CAR cells. The trajectory complexity was reduced by excluding a branch that mapped to chondro-primed OLCs based on label transfer.

We performed receptor perturbation simulations on the subclustered MSC data using IntraTalker. We defined candidate receptors as those connected to downstream transcription factor targets within the for MSCs predicted receptome. Further, we performed perturbation analysis separately on each condition (KO, WT). We trained ridge regression models (*α* = 10) using gene expression and receptome information to obtain a coefficient matrix for each condition. The perturbation process was performed over five iterations. The resulting shifts were projected onto the trajectory embedding as a vector field. We quantified changes in pseudotime and assessed statistical significance using the Wilcoxon signed-rank test. We calculated effect sizes using Cohen’s d and ranked receptors accordingly.

### Preprocessing and Cell–Cell Communication Analysis of Myocardial Infarction snRNA-seq Data

We retrieved the AnnData^62^ object of the snRNA-seq data from Kuppe et al.^24^. This included raw counts, along with sample annotations and two cluster level (main and sub-cluster levels). We further updated the sub-cluster annotation names to improve interpretability, particularly for cardiomyocytes and fibroblasts (Table S3). Further, we removed cells annotated as proliferating to minimize cell cycle-related transcriptional bias in the cell-cell communication analysis. We normalized the data with Scanpy (v1.11.5)^64^ using default parameters and filtered genes to be present in at least 1500 cells.

We inferred ligand-receptor interactions using CellphoneDB v2.0^63^ via the LIANA+ framework (v1.6.1)^11^ and the human consensus database^11^, with the LR expression proportion threshold set to 0.1. Predicted ligand-receptor interactions with *p* < 0.05 were considered for downstream analysis. For transcription factor activity analysis, we included the gene regulatory networks inferred from snATAC-seq with scMega^16^ provided in Kuppe et al.^24^ for cardiomyocytes and fibroblasts. We used IntraTalker to merge the DoRothEA^25^ regulome with confidence levels A and B with the gene regulatory networks. To emphasize the transcription factor-target gene interactions from the gene regulatory, the weights of DoRothEA database interactions were reduced from 1.0*/* −1.0 to 0.5*/* −0.5, and from 0.5*/* −0.5 to 0.25*/* −0.25. This resulted in cardiomyocyte- and myofibroblast-specific regulomes.

Transcription factor activity and IntraTalker analyses were performed separately for cardiomyocytes and fibroblasts using the corresponding cell-type-specific regulomes. In parallel, analysis was performed on all remaining cell types using only the DoRothEA regulome with confidence levels A and B. Transcription factor activities were estimated using the decoupler (v2.1.2)^53^ ULM method. IntraTalker was run in all cases with a p-value threshold < 0.05 and a log fold-change threshold > 1. The results of the intracellular signalling analysis from cardiomyocytes, fibroblasts, and the other cell types were first combined and then integrated with the predicted ligand-receptor interactions. Then we performed differential cell-cell communication analysis with CrossTalkeR2 using IntraTalkers receptor perturbation module and the receptome predicted from the transcription factor activity analysis, we ran receptor stimulation simulations using all receptors with downstream signalling.

Models were trained using standard parameters and perturbations were run with five iterations. We ranked receptors by the changes they induced with IntraTalkers standard processing. For the downstream ligands of the top-ranking receptors, we performed enrichment analysis with clusterProfiler (v4.20.0)^67–69^ against the Gene Ontology Biological Process terms (GO.db v3.23.1)^43,44^. As the background (universe) gene set, we used all genes expressed in at least 1% of cells (expression fraction ≥ 0.01).

### Marker Selection in Myocardial Infarction Case Study for qPCR Experiments

We identified marker genes associated with SCARA5^+^ fibroblasts and myofibroblasts along the pseudotime trajectory using Spearman correlation (SciPy v1.16.2) in Python and generalized additive models implemented in the tradeSeq package (v1.26.0) in R^46^. For tradeSeq, models were fitted using six knots, and the startVsEndTest() function was applied to identify genes differentially expressed between the beginning and end of the trajectory and to obtain Wald statistics. Marker genes were selected by combining Spearman correlation coefficients with tradeSeq Wald statistics, prioritizing genes that showed both strong correlation with pseudotime and significant differential expression along the trajectory.

### Normal human ventricular cardiac fibroblasts (NHCF-V) Culture

NHCF-V (Lonza, CC-2904) were cultured in FGM™-3 Cardiac Fibroblast Growth Medium-3 BulletKit™ (Lonza, CC-4526). NHCF-V were maintained in a humidified incubator at 37°C with 5% CO_2_. For subculture, NHCF-V at 80% confluence was washed with DPBS (Bio&SELL, L1835), detached using Accutase Solution (PromoCell, C-41310), and neutralized with DPBS. After centrifugation at 200 ×g for 5 min, NHCF-V were seeded at a density of 3,500 cells/cm^2^. The culture medium was changed every 2 days. NHCF-V between passages 3 and 5 were used for subsequent experiments.

NHCF-V were seeded into 6-well plates. When cell confluence reached 80%, the culture medium switched to FGM−-3 containing 1% FCS for overnight starvation, the following day, treatment was initiated using 50 ng/mL IGF-1 [stock solution, 50 µg/mL], (Thermo Scientific, 100-11-100µg), 10 ng/mL TGF-*β* 1 [stock solution, 10 µg/mL], (Thermo Scientific, 100-21-10µg). The medium contained 0.1% BSA at a dilution of 1:1000, used as the control group. The culture medium was changed every 2 days. 96 h after treatment, the NHCF-V were harvested for RNA extraction.

### Quantitative Real-Time PCR (qPCR)

Total RNA was extracted using the RNeasy Mini Kit (QIAGEN, 74104) according to the manufacturer’s protocol. RNA was eluted with RNase-free water, and its concentration and purity were determined using a NanoDrop Lite (Thermo Scientific). Only RNA samples with an A260/A280 ratio between 1.8 and 2.0 were used for subsequent experiments. For each reaction, 2µg of total RNA was used to synthesize complementary DNA (cDNA) using the High Capacity cDNA Reverse Transcription Kit (Thermo Scientific, 4368813), following the manufacturer’s protocol. The synthesized cDNA was subsequently used for subsequent experiments.

Quantitative Real-Time PCR (qPCR) was performed in a 10µL reaction mixture containing 10 ng cDNA, specific primers, and 2× primaQUANT ADVANCED qPCR Master Mix (Steinbrenner Laborsysteme GmbH, SL-9903). The amplification was performed following the manufacturer’s standard 3-step protocol. Data were analyzed using the 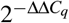method and normalized to the reference gene GAPDH. Primers for genes are listed in the Supplementary table S4.

### Statistical Analysis

For qPCR, relative gene expression was compared using a linear mixed-effect model implemented in the Python library *statsmodels* v0.14.6 to account for the non-independence of measurements from biological and technical replicates. For each gene, we consider the following model :

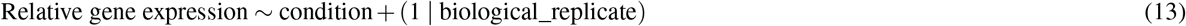

where condition (BSA, IGF1, TGFB1) was modeled as a fixed effect, and biological replicate was included as a random intercept, allowing the three technical replicates measured within each biological sample to share a common baseline rather than being treated as independent observations. Pairwise comparisons of treatment conditions against the BSA control (IGF1 vs. BSA; TGFB1 vs. BSA) were extracted from the fitted model, and p-values were corrected for multiple comparisons using the Holm-Bonferroni method.

## Data and Code availability

The Code, tutorials, and examples that replicate the analysis in this manuscript are available at https://github.com/CostaLab/IntraTalker, https://github.com/CostaLab/IntraTalkerpy, https://github.com/CostaLab/CrossTalkeR and https://github.com/CostaLab/pyCrossTalkeR. Pre-processed objects are found in Zenodo https://doi.org/10.5281/zenodo.20813792.

## Acknowledgements

This project has been funded by the Federal Ministry of Research, Technology and Space of Germany (BMFTR) via the Consortia E:MED Fibromap and CureFib.

## Author information

V.K., J. N. and I.C. conceived the experiment(s), V.K. conducted the experiment(s), V.K and I.C. analyses the results. J.N., L.H., J. M. and M.R. supported software and method developments. Z.F. and C.K. planned and performed the *in vitro* experiments. All authors reviewed the manuscript.

## Supplementary

**Figure S1.**
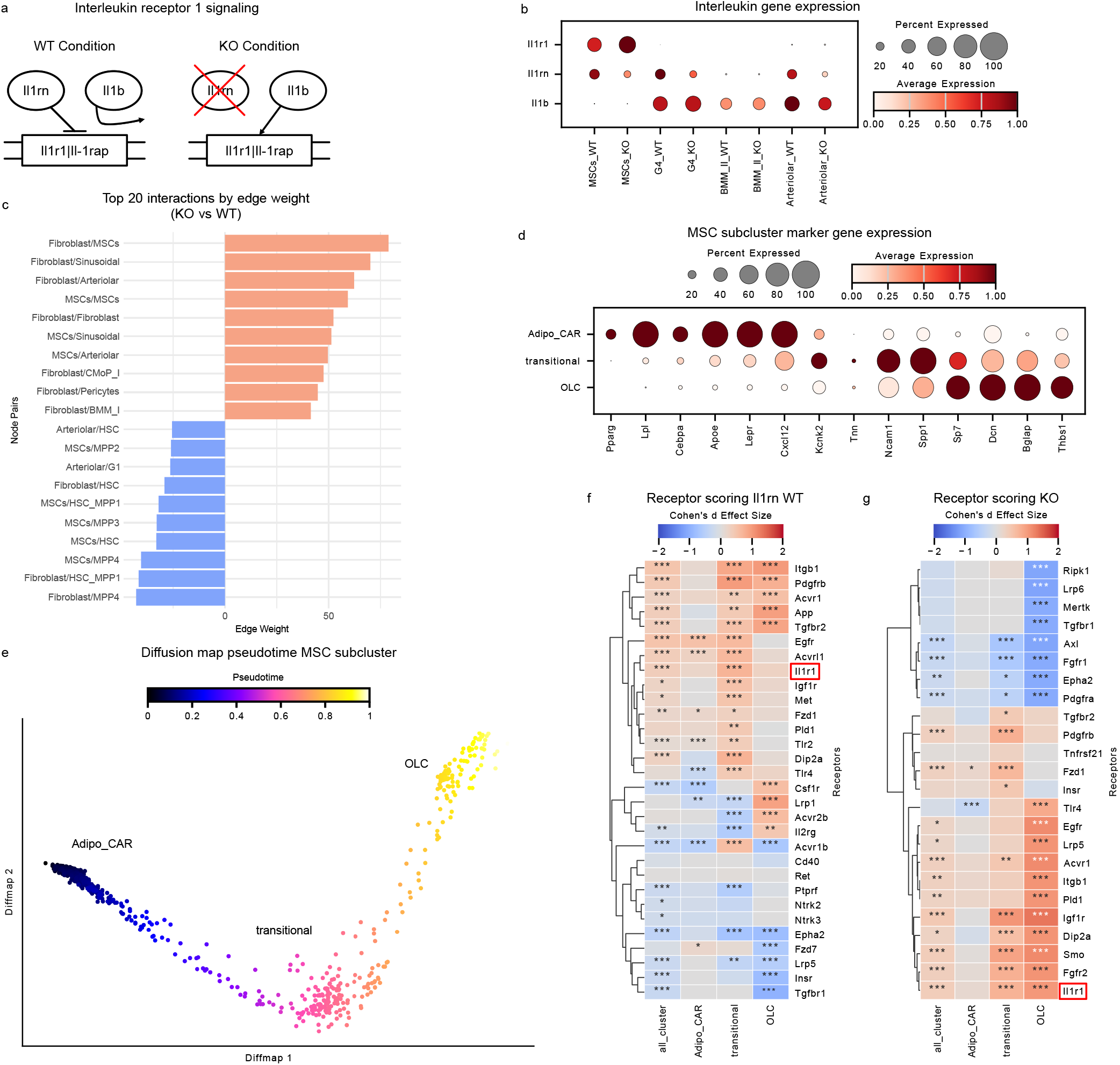
**a** Experimental set up for the interleukin signalling study. In WT condition the receptor complex Il1r1-Il1rap is blocked by Il1rn. In the Il1rn KO condition Il1b can bind to the complex and interleukin signalling via Il1b is activated. **b** Dotplot showing the expression of the interleukin signalling genes Il1r1, Il1rn and Il1b for selected cells in WT and KO conditions. **c** Top 20 cell-cell interactions by signalling weight comparing KO and WT conditions. **d** Dot plot showing marker genes within the MSC subclusters. **e** Diffusion map representation of MSCs sub-clusters (AdipoCAR, transitional and OLCs, n=828). The color indicates the pseudo-time, which increases from AdipoCAR towards OLCs. **f**,**g** Receptor scores after perturbation experiments in WT condition ( **f**) and Il1rn KO condition (**g**). Color scale reflects effect size score (Cohen’s d) based on perturbation-induced difference in pseudotime. Receptors are clustered by score similarity. \**P* < 0.05, ** *P* < 0.01, \*\*\**P* < 0.001 (Wilcoxon signed-rank test). We highlight the enrichment of Il1r1 in WT and KO conditions.

**Figure S2.**
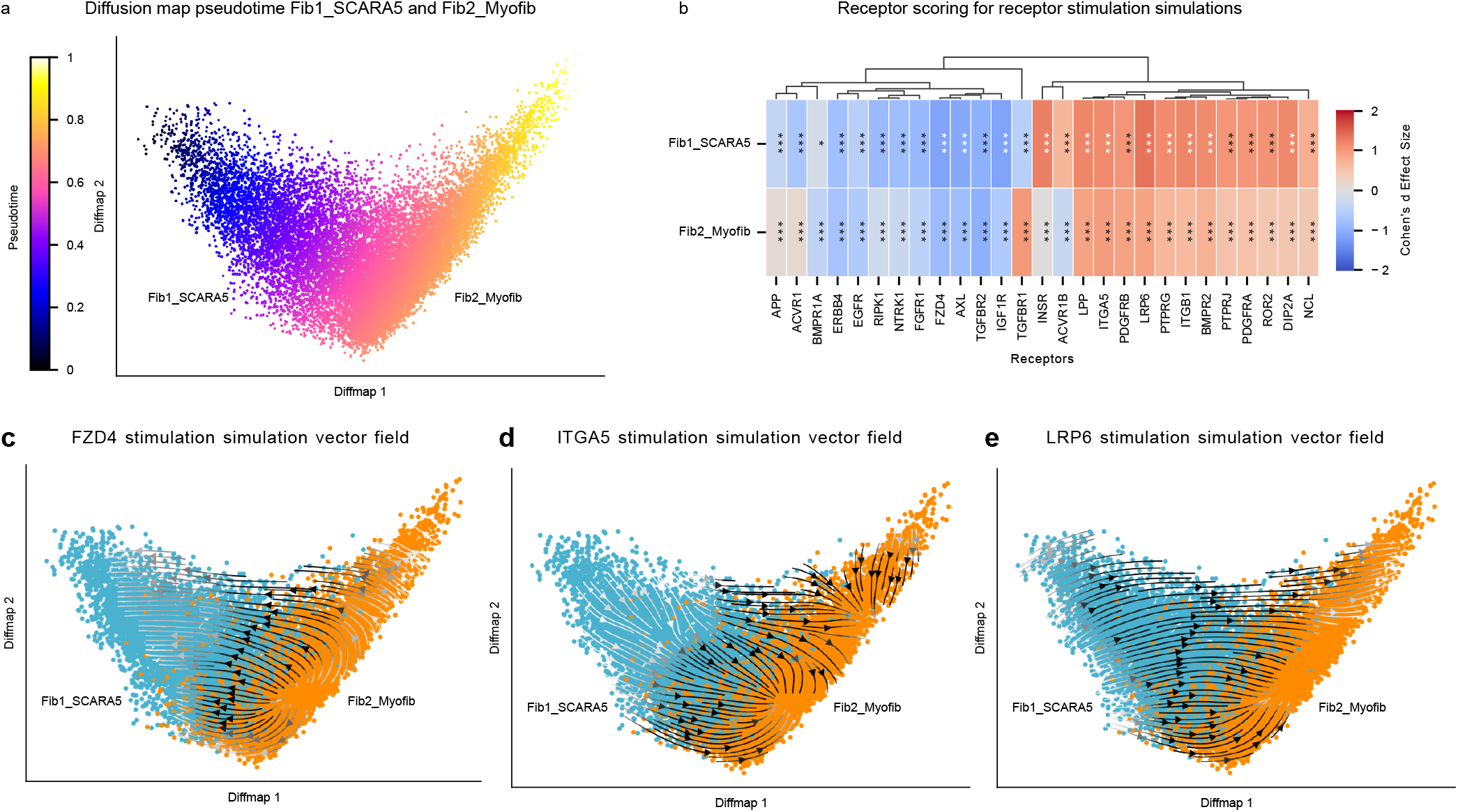
**a** Diffusion map representation of SCARA5^+^ fibroblasts and myofibroblasts. The color indicates the pseudo-time, which increases from SCARA5^+^ fibroblasts towards myofibroblasts. **b** Receptor scores after perturbation experiments in all samples. Color scale reflects effect size score (Cohen’s d) based on perturbation-induced difference in pseudotime. Receptors are clustered by score similarity. \**P* < 0.05, ** *P* < 0.01, *** *P* < 0.001 (Wilcoxon signed-rank test). **c** Predicted shifts after FZD4 stimulation simulation in SCARA5^+^ fibroblasts and myofibroblasts. Arrows are shaded by the distance each cells shifted after receptor simulation. **d-e** Same as c for ITGA5 and LRP6.

**Figure S3.**
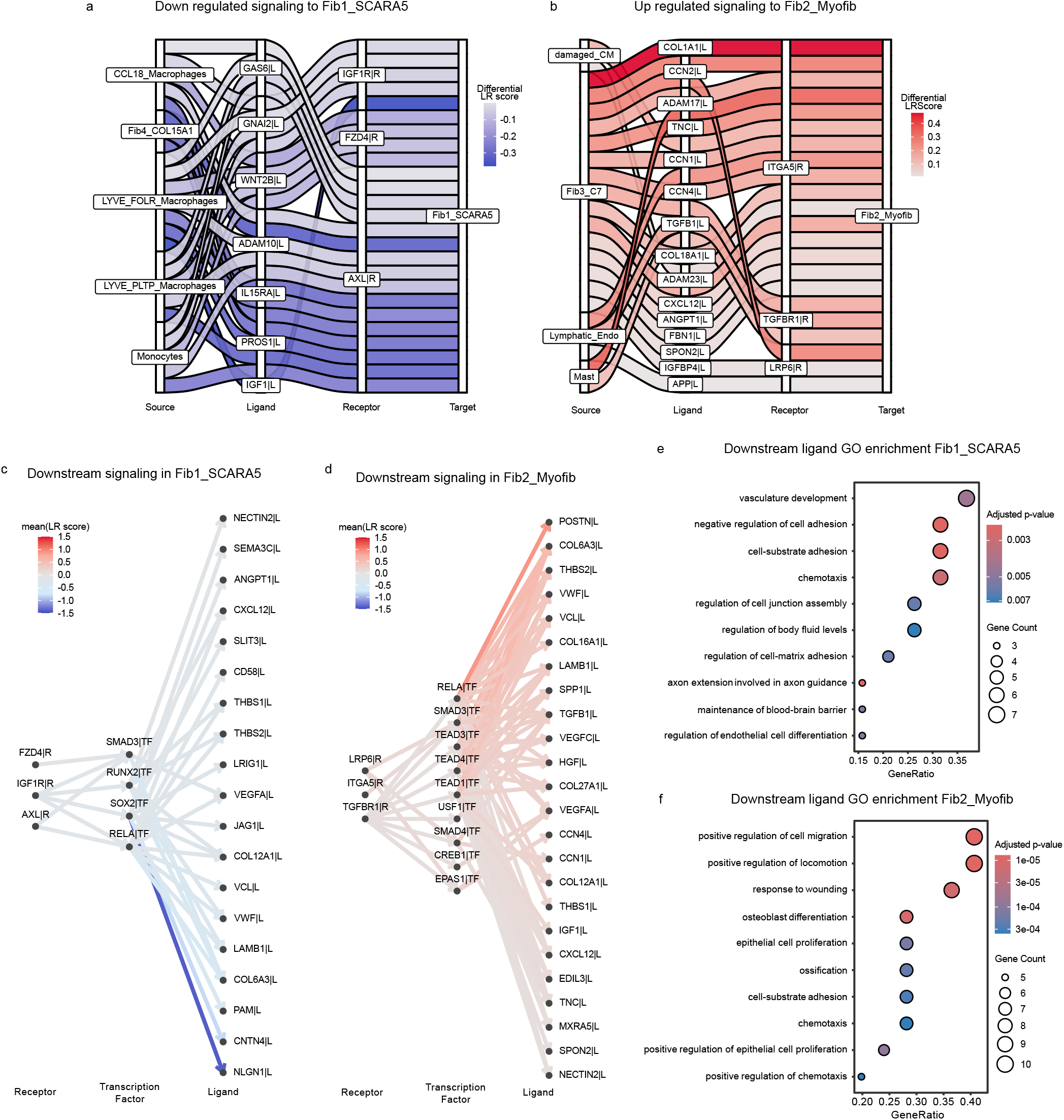
**a**,**b** Sankey plot of interactions associated with the myogenic condition in the myeloid-fibroblast community towards SCARA5^+^ fibroblasts (Fib1_SCARA5) (a) and interactions associated with the ischemic condition in the damaged-fibrotic community towards myofibroblasts (Fib2_Myofib) (b) for selected receptors. The edge colors indicate the differential LR Score. **c**,**d**. Downstream signalling of selected receptors in Fib1_SCARA5 cells associated with the myogenic condition (c) and in Fib2_Myofib cells associated with the ischemic condition. The color scale indicates the relative mean LR score of the incoming receptor and outgoing ligands. **e**,**f** GO enrichment results performed on target ligands from **c** and **d** for Fib1_SCARA5 (e) and Fib2_Myofib (f) cells. *P* values were adjusted for multiple comparisons using the Benjamini-Hochberg correction.

## 1 Related Work

Over the past years, many tools for inferring cell-cell communication have been introduced, primarily focusing on ligand-receptor interactions derived from transcriptomic data^7^. More recent methods extend this approach by incorporating intracellular downstream signalling as a consequence of intercellular communication. This is typically modeled through transcription factor activity or pathway activity inference, using either correlation-based approaches or prior-knowledge networks such as protein-protein interactions and gene regulatory networks. Here, we focus on methods that leverage prior knowledge to infer intracellular signalling while constructing cell-cell communication networks, including NicheNet^70^, scMLnet^8^, scSeqComm^9^, CellphoneDB (v4.1)^10^, LIANA+^11^, and IntraTalker. Most of these methods utilize transcription factor activity analysis to model intracellular signalling, differing primarily in the statistical method used to infer transcription factor activity and how signalling is propagated.

At gene-level resolution, multilayer network approaches such as scMLnet^8^ construct downstream signalling by connecting ligands, receptors, transcription factors, and target genes. Intracellular signalling is inferred by linking receptors to transcription factors via interactions and pathway networks, and transcription factors to target genes via gene regulatory networks. Receptor-transcription factor links and transcription factor-target gene activity are approximated using Fisher’s exact tests and further filtered by expression correlation on interacting partners. The framework enables gene-level resolution of signalling cascades from ligands to downstream targets. However, the proxy measures of TF activity limit quantitative interpretability. Additionally, the network is unweighted and does not support differential or condition-specific analysis. Lastly, the multilayer networks for each cell type are disconnected and do not allow propagation of downstream activation of ligands towards other cells.

Complementing gene-level resolution approaches, scSeqComm^9^ models signalling through ligand-receptor-transcription factor relationships but focuses on functional characterization at the pathway level rather than at the individual gene level. As with scMLnet, transcription factor activities are inferred using Fisher’s exact test and additional differential gene expression analysis. The more recent extension, scSeqCommDiff^71^, allows differential analysis of samples and conditions.

In contrast to scMLnet and scSeqComm, CellphoneDB and LIANA+ apply dedicated methods to infer single-cell transcription factor activities. CellphoneDB (v4.1)^10^ extends the original ligand-receptor framework by incorporating downstream signalling via transcription factor activity inference with VIPER and the DoRothEA regulon database. Transcription Factor Activity is estimated from the enrichment of regulon targets in ranked gene expression signatures, enabling quantitative, cell-specific estimates of transcription factor activity. It also introduces a collection of well-described receptor-transcription factor links. However, the scope of the intracellular signalling is limited. It does not explicitly construct receptor-to-target signalling paths and also does not propagate signals beyond target genes.

Similarly, LIANA+^11^ provides a flexible framework that unifies multiple ligand-receptor inference methods with transcription factor activity analysis with decoupler^53^. It employs linear models such as the univariate linear model (ULM) to estimate transcription factor activity from gene expression data. It further connects ligand-receptor interactions with downstream transcription factors, utilizing prior knowledge from pathway databases. However, it does not explicitly model downstream target-gene regulation and thus does not enable analysis beyond transcription factors or target genes.

A conceptually distinct strategy is presented by NicheNet^70^, which does not rely on transcription factor activity analysis to predict intracellular downstream signalling. Instead. NicheNet prioritizes ligand-target gene relationships using prior-knowledge-based signalling and gene regulatory networks. Ligands of interest are defined based on gene expression, and their downstream effects are inferred, linking them to differentially expressed genes in the receiver cell. The extension MultiNicheNet^72^ enables differential cell-cell communication analysis of multi-sample, multi-condition single-cell data. While NicheNet provides gene-level associations between ligands and targets, it does not explicitly construct signalling graphs across distinct cells and is strongly dependent on the accuracy of prior knowledge.

Intra Talker expands all previous approaches by combining transcription factor inference using decoupler with receptor-centered signalling networks that connect receptors to downstream target genes. As scMLnet, IntraTalker reconstructs signalling from ligands to target gene at gene-level resolution, but by the adaptation of the CrossTalkeR2 framework, it allows propagation of signals beyond one cell and supports multi-condition analysis. This allows for signalling effects to be linked into a broader communication context.

A recent tool, CellAgentChat^73^, transfers the concept of transcription factor expression perturbation directly into the context of cell-cell communication by using receptor expression perturbations. The goal is to prioritize receptors based on the perturbation-induced changes and their importance in disease conditions. For the latter, CellAgentChat uses survival analysis and the database DisGeNET to connect perturbed genes to the disease state of interest. The tool does not perform transitional modelling to infer changes in cell states, nor does it consider inferred gene regulatory networks, but instead relies on prior knowledge signalling databases for modelling. In contrast, IntraTalker performs receptor perturbation in the context of receptor-to-target signalling networks and relates predicted transcriptional changes to cellular trajectories, enabling the assessment of receptor impact on cell state transitions.

Overall, current approaches address different aspects of intracellular signalling, including transcription factor activity inference, receptor-to-target modelling, and perturbation analysis, but mostly do not integrate intracellular and intercellular components (see Sup. Table S1). IntraTalker unifies these aspects by combining gene-level transcription factor activity inference with receptor-centered signalling networks and receptor perturbation analysis. In addition, its integration with CrossTalkeR2 enables differential analysis of inter- and intracellular communication across conditions, extending beyond existing approaches, where multi-condition analysis is limited.

**Table S1.** Characteristics of Methods associated with Related Work.

| Tool | TF Activity Score | Intracellular signalling (Gene-Level) | R-TF Weighting | Cross-Cell Interactions | GRN- based | ATAC-seq GRN | Multiple Conditions | Receptor Expression Perturbation | Receptor Prioritization | Transitional Modelling |
| --- | --- | --- | --- | --- | --- | --- | --- | --- | --- | --- |
| NicheNet (+MultiNicheNet) | x | o | ✓(a priori) | x | ✓ | x | ✓ | x | x | x |
| scMLnet | o | ✓ | ✓ | x | ✓ | x | x | x | x | x |
| scSeqComm (+scSeqCommDiff) | o | ✓ | ✓(a priori) | x | ✓ | x | ✓ | x | x | x |
| LIANA+ | ✓ | ✓ | ✓ | x | ✓ | x | x | x | x | x |
| CellphoneDB | ✓ | ✓ | x | x | ✓ | x | x | x | x | x |
| CellAgentChat | x | x | x | x | ✓ | x | x | ✓ | ✓ | x |
| IntraTalker+CrossTalker2 | ✓ | ✓ | x | ✓ | ✓ | ✓ | ✓ | ✓ | ✓ | ✓ |

**Table S2.**
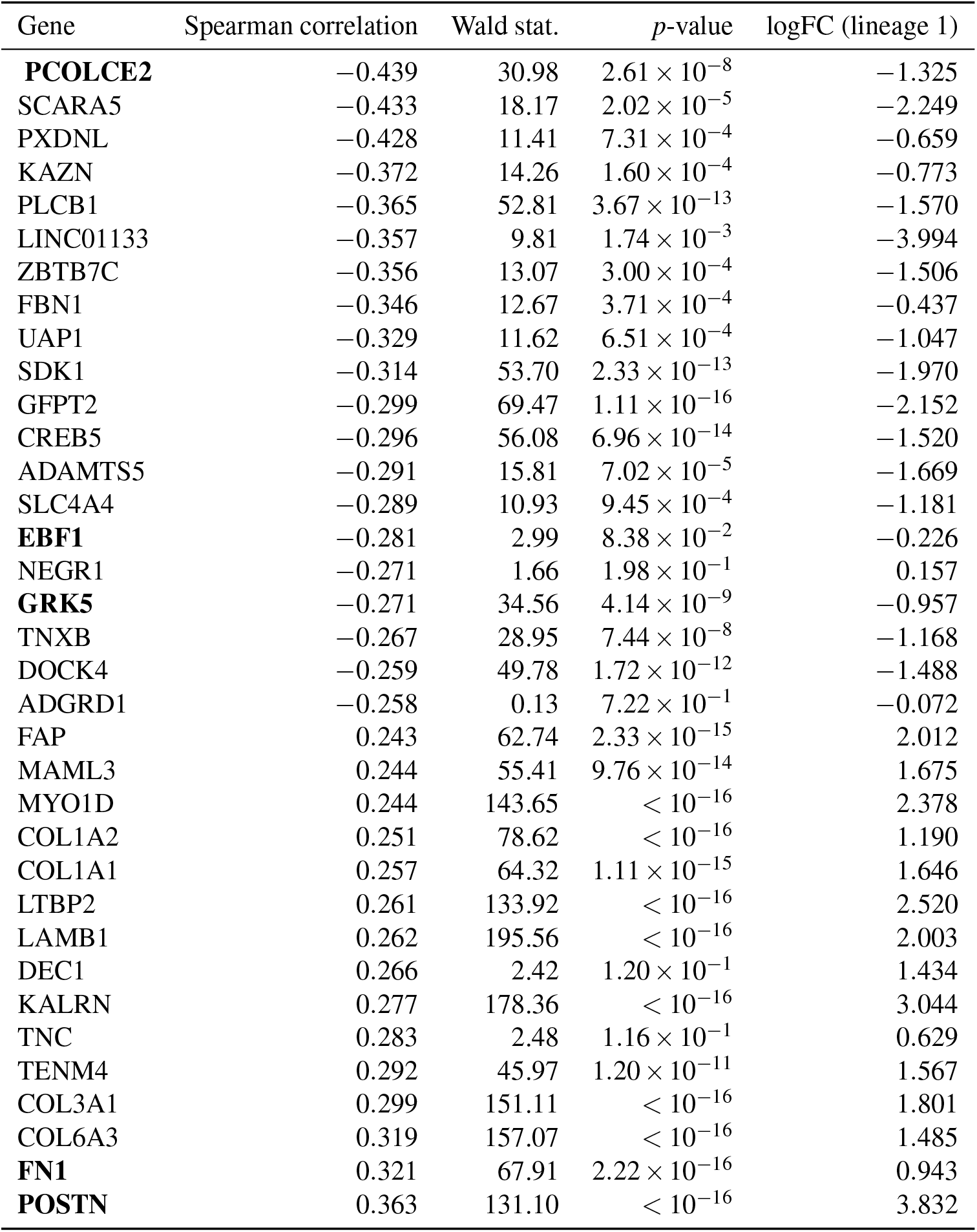
Marker genes along the SCARA5^+^ fibroblast-to-myofibroblast pseudotime trajectory.

**Table S3.** Updated Sub-Cluster Annotations for the Myocardial Infarction Heart Data.

| Original Annotation | Updated Annotation |
| --- | --- |
| Fib1 | Fib1_SCARA5 |
| Fib2 | Fib2_Myofib |
| Fib3 | Fib3_C7 |
| Fib4 | Fib4_COL15A1 |
| vCM1 | healthy_CM |
| vCM2 | intermediate_CM |
| vCM3 | damaged_CM |
| vCM4 | vCM3 |
| vCM5 | vCM4 |

**Table S4.**
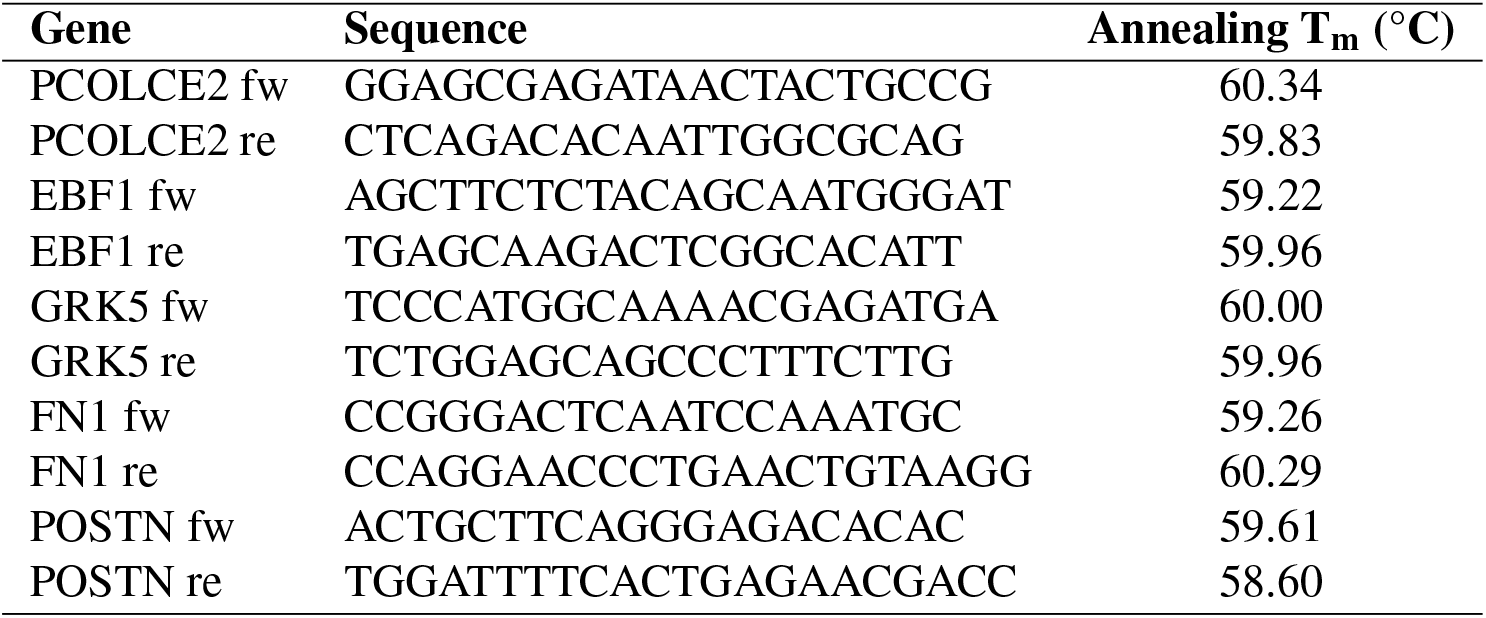
Primer sequences and annealing temperatures.

